# Development and validation of PainFace, a software platform that simplifies and standardizes mouse grimace analyses

**DOI:** 10.1101/2022.08.12.503790

**Authors:** Eric S. McCoy, Sang Kyoon Park, Rahul P. Patel, Dan F. Ryan, Zachary J. Mullen, Jacob J. Nesbitt, Josh E. Lopez, Bonnie Taylor-Blake, James L. Krantz, Wenxin Hu, Rosanna L. Garris, Lucas V. Lima, Susana G. Sotocinal, Jean-Sebastien Austin, Adam D. Kashlan, Jessica A. Jimenez, Sanya Shah, Abigail K. Trocinski, Kelly A. Vanden, Rami M. Major, Hannah O. Bazick, Morgan R. Klein, Jeffrey S. Mogil, Guorong Wu, Mark J. Zylka

**Author notes:** equally-contributed (Eric S. McCoy, Sang Kyoon Park, Rahul P. Patel).

## Abstract

Facial grimaces are now commonly used to quantify spontaneous pain in mice and other mammalian species, but scoring remains subjective and relies on humans with very different levels of proficiency. Here, we developed a Mouse Grimace Scale (MGS) for black-coated (C57BL/6) mice consisting of four facial action units (orbitals, nose, ears, whiskers). We used this scale to generate ground truth data from over 70,000 images of black mice in different settings. With this large data set, we developed a deep neural network and cloud-based software platform called PainFace (http://painface.net) that accurately scores facial grimaces of black mice on a 0–8 scale. PainFace generates over two orders of magnitude more MGS data than humans can realistically achieve, and at superhuman speed. By analyzing the frequency distribution of grimace scores, we found that mice spent >7x more time in a high grimace state following laparotomy surgery relative to sham surgery controls. The analgesic carprofen reduced the amount of time animals spent in this high grimace state after surgery. Specific facial action unit score combinations were overrepresented following laparotomy surgery, suggesting that characteristic facial expressions are associated with a high grimace state. We performed validation experiments in two labs located in different countries to demonstrate reproducibility of the PainFace platform. To further enhance rigor and reproducibility, we will invite pain researchers to beta test PainFace and then incorporate their feedback into the software and manuscript prior to peer review. While this study is focused on mice, PainFace was designed to simplify, standardize, and scale up grimace analyses with many other mammalian species, including humans.

## Introduction

Pain in rodents has traditionally been measured using evoked behavioral responses to experimenter-delivered nociceptive stimuli, such as paw withdrawal from a noxious heat stimulus or nylon monofilament. However, such evoked responses do not properly model spontaneous, non-evoked pain—the primary symptom experienced by chronic pain patients (Backonja, 2012; Maier et al., 2010). Facial grimaces can be used to quantify spontaneous pain in mice and other mammalian species (Dalla Costa et al., 2016; Di Giminiani et al., 2016; Gleerup et al., 2015; Guesgen et al., 2016; Häger et al., 2017; Holden et al., 2014; Keating et al., 2012; Matsumiya et al., 2012; Mogil et al., 2020; Reijgwart et al., 2017; van Loon and Van Dierendonck, 2015). The original Mouse Grimace Scale (MGS) was developed for white-coated mice and has been used to evaluate responses to noxious stimuli and the efficacy of analgesics (Langford et al., 2010). While the MGS has been used by many labs, manual scoring of facial grimacing is labor intensive and humans vary in their scoring proficiency.

The manual MGS scoring process entails collecting high-quality videos of mice with their faces clearly in view. Then a relatively small number of frames, typically 8-10, from each video are extracted and scored by experts. These experts identify specific “facial action units” (FAUs) such as orbital tightening (closed eyes), and then must assess whether the action unit is absent/not present (score of 0), moderate (score of 1), or obvious (score of 2). Scores for each action unit can be averaged, as described (Langford et al., 2010), or summed (as done in our present study) to calculate the MGS score for one video frame.

Partial automation was achieved via the development of Rodent Face Finder software, which generated human-scorable picture files from videos (Sotocinal et al., 2011). To further automate this process and standardize scoring, we developed a machine-learning algorithm that categorized individual images of white mice into binary low (“no pain”) or high (“pain”) grimace categories (Tuttle et al., 2018). However, the algorithm does not output numerical MGS scores nor does it work with black-coated C57BL/6 mice, the mouse strain most commonly used by neuroscientists and pain researchers (Sadler et al., 2022). Moreover, the software has to be downloaded, installed, and operated using a command line interface, which is not intuitive for most researchers to use.

Here, we sought to overcome these and other barriers that have limited the widespread use of facial grimaces to quantify spontaneous pain. In pursuit of this goal, we developed a user-friendly software platform called PainFace that simplifies and standardizes the quantification of facial grimaces from black-coated mice. PainFace utilizes deep neural networks to identify and score facial action units. While our study is focused on black-coated mice, PainFace has broad future utility. The machine learning backbone and software were designed to facilitate grimace analyses with mice of any coat color and other mammalian species.

## Materials and Methods

### Animal care and use

All procedures used in this study were approved by the Institutional Animal Care and Use Committee at the University of North Carolina at Chapel Hill or the University Animal Care Committee at McGill University. At UNC, group-housed C57BL/6 mice (The Jackson Laboratory, 8-12 weeks of age, sex indicated in figure legends) were maintained on a 12:12 h light/dark cycle and given food (PicoLab 5V5R, LabDiet) and tap water *ad libitum*. At McGill University, group-housed C57BL/6 mice (The Jackson Laboratory, 6-8 weeks of age) were maintained on a 12:12 h light:dark cycle and given food (Teklad 8604, Envigo) and tap water *ad libitum*.

### Imaging setups

Over the course of this study, two different imaging setups were used to collect videos of black-coated mice. Our original mouse behavioral chamber was made of ¼” stainless steel with a clear ½” Plexiglas back; internal dimensions: 5.5 cm width, 14 cm depth, including the 4-cm extension beyond the floor, 6 cm height from the floor to the top of the chamber. The key light (5500K LEDs, 600 LED Video Studio Lighting Photography Panel, ePhoto Inc.) was placed above the chamber and angled towards the front. One Canon Vixia HF R800 camera was placed 15-20 cm from the chamber opening. The light level was increased until the mouse face was clearly visible and brightly lit with no shadows. Once the chamber was illuminated, exposure was increased by +0.75. Auto focus was set to ON. Program was set to AE so the camera controls shutter speed and aperture. Zoom speed was set to VAR. Framing Assistance was set to AUTO. Focus Assistance, Face Detection & Tracking, Auto Backlight Correction, Slow Shutter Speed were set to ON. Flicker Reduction was set to OFF. Powered IS was OFF. White Balance was set to AWB.

Our optimized imaging system included a custom-designed behavioral chamber made from aluminum, stainless steel, and opaque plastic (http://hypothesistohardware.com, Mouse Behavioral Chamber). The chamber was mounted to an aluminum breadboard (ThorLabs, # MBH1224). Top lights (4000K, 100% intensity, Viltrox 50W/4700LM 3300K-5600K bi-color) and key lights (5600K, 100% intensity, Neewer Advanced 2.4G 480 LED) were used for illumination. Arducams (12.3 MP 1/2.3 inch IMX477 camera module with 8 mm CS-Mount Lens, metal enclosure) were mounted at a fixed distance from the behavioral chamber, and videos were stored on Raspberry Pi computers (CanaKit Raspberry Pi 4 4GB RAM) prior to transferring to PainFace. Camera and lights were affixed to the breadboard using optomechanical parts (ThorLabs, TR12, TR4, TR1.5, RA90, and TR6).

### Software development

PainFace is an integrated cloud-based platform for managing, analyzing, and visualizing grimace data worldwide. PainFace consists of a Restful API server, a web-based user interface, and workers (see below) that evaluate grimace videos. Use of the industry-standard REST (<u>Re</u>presentational <u>S</u>tate <u>T</u>ransfer) API enables external users and researchers to develop their own software, such as custom user interfaces or automation scripts, to access PainFace workers (Fielding, 2000). The API server also has login and user management functions to enable research groups to manage their resources separately and securely.

The web-based user interface can be used to annotate datasets and evaluate videos with various machine learning models. Mouse strain, species, sex, and other key experimental parameters can be designated with corresponding metadata. Users can select pre-built detection models to locate bounding boxes around action units and predict grimace scores for each facial action unit.

PainFace permissions are separated into three levels: standard user, scorer/annotator, and supervisor. Standard users can upload videos, initiate grimace analyses, and view analysis results—the main functions needed by the vast majority of all researchers. Scorer/annotators have access to tools for generating ground truth data—functions to extract images from videos and manually score and annotate the images. Supervisors can manage all video files, including uploading and deleting videos.

Workers are scalable, independent modules that identify features and generate grimace scores for each action unit. We used standard messaging queue technology to schedule the real-time computation workload. This distributed scheduling makes it possible to scale and process large numbers of videos in parallel. Workers can process up to 32 full high-definition videos (1920 x 1080 RGB, 30 frames per second; fps) simultaneously on one node. One computation node consists of one 8-core CPU, 128 GB RAM, and two Nvidia RTX 3090 graphic cards (with 24 GB GPU memory and 10,496 GPU cores).

### Laparotomy surgeries

All experiments were performed during the light phase at approximately the same time each morning. Prior to placing mice in the imaging setup, the mice were acclimated to the testing room for 30 minutes in their home cages. Animals were then placed into the imaging chamber. A 30-min baseline video was acquired at 30 fps. Mice were then removed from the chamber and were anesthetized using 5% isoflurane until hindpaw areflexia. Mice were maintained at 1.75% isoflurane for the remainder of the surgery. Eyes were lubricated with Blink^®^ Tears drops. We recommend against the use of petroleum-based eye lubricants because they tend to cake on the eyes, causing eyes to remain closed. The abdomen was sprayed generously with ethanol. A small area was shaved using a razor blade between the ribs and hips, avoiding nicking the skin, especially of sham surgery animals, as tissue damage elsewhere has the potential to cause grimacing independent of the laparotomy surgery. The area was cleaned three times, alternating between betadine and chlorhexidine. The area was then sprayed once more with ethanol to remove the excess betadine and chlorhexidine. Excess ethanol was wiped away using a sterile cotton pad. A 1.5-cm transverse incision was made to the outer skin layer just above the hips in the center of the abdomen. A 1-cm incision was made to the peritoneum to expose the intestines. Approximately 1-2 cm of intestines was removed from the cavity and palpated for 10 s using forceps. The intestines were returned to the abdominal cavity, and the peritoneum was sutured. Tissue adhesive was used to close the outer incision. The entire body of the animal was then wiped down with a sterile cotton pad to remove any excess fluid from the fur, so that the fur does not look “spikey” in the videos. Mice were placed supine in a cage and heated on a warming pad for 30 min to facilitate recovery. After the 30-min recovery period, the mice were returned to the imaging chamber and recorded for an additional 30 min.

### Validation experiment–UNC Chapel Hill

Mice were separated into three groups: sham, laparotomy (LAP), and carprofen + laparotomy (CARLAP). Sham mice were anesthetized and treated identical to LAP and CARLAP mice, including shaving of the fur, but no incision was made. Carprofen (Rimadyl, 50 mg/kg s.c., Med Vet International, #RXRIM-INJ, starting conc. 50 mg/mL) was administered immediately before shaving.

### Validation experiment–McGill University

For carrageenan experiments (no habituation), 10 C57BL/6 male mice (6 weeks of age) were acclimated to the recording chamber for 30 min with lights on. After acclimation, the mice were recorded for a 30-min baseline video. Next, the mice were bilaterally injected into the hindpaw with 20 µL of 1% carrageenan and an additional recording for 1 hour was taken 3 hours post-injection. For the carrageenan habituation experiments, 10 C57BL/6 male mice were habituated in the recording chamber with the lights on for 1 hour each day for 3 days prior to administration of carrageenan.

### Inter-rater reliability assessment

Twenty-five frames were randomly selected and independently scored by four trained scorers (Zylka lab n=3 scorers, Mogil lab n=1 scorer). Scoring reliability was determined by Intraclass Correlation coefficient (two-way mixed design) in SPSSv28.

### Statistics

All statistical analyses were performed in GraphPad Prism or SPSS v.28.

## Results

### Optimized imaging conditions for black-coated mice

We initially encountered problems while generating training data with black-coated mice. The ambient room lighting conditions that we previously used to visualize the facial action units of white-coated mice were too dark to visualize most of the facial action units of black-coated mice (Tuttle et al., 2018). We were unable to consistently find the eyes against the black fur, let alone determine if the eyes were open or closed, nor were we able to consistently identify the nose, the cheeks, or the whiskers (Supplementary Figure 1A). We found that supplemental lighting was needed to clearly see the face of black-coated mice (Supplementary Figure 1B). To facilitate reproducible placement of the lights, we built an imaging setup from commercially available optomechanical parts, LED lights, and a high-definition (1920 x 1080) color video camera. We also designed a behavioral imaging chamber that can be mounted to a metal breadboard at a fixed distance from the lights and camera (Figure 1A). This optimized behavioral chamber includes an adjustable rear wall to accommodate mice of different sizes, a removable floor that is easy to clean and can be slid into the chamber along with the mouse, and an open front with elevated cliff, which encourages mice to face the camera (Tuttle et al., 2018). We recommend that imaging conditions similar to those shown in Figure 1B be used when acquiring videos of black-coated mice for grimace analyses.

**Figure 1.**
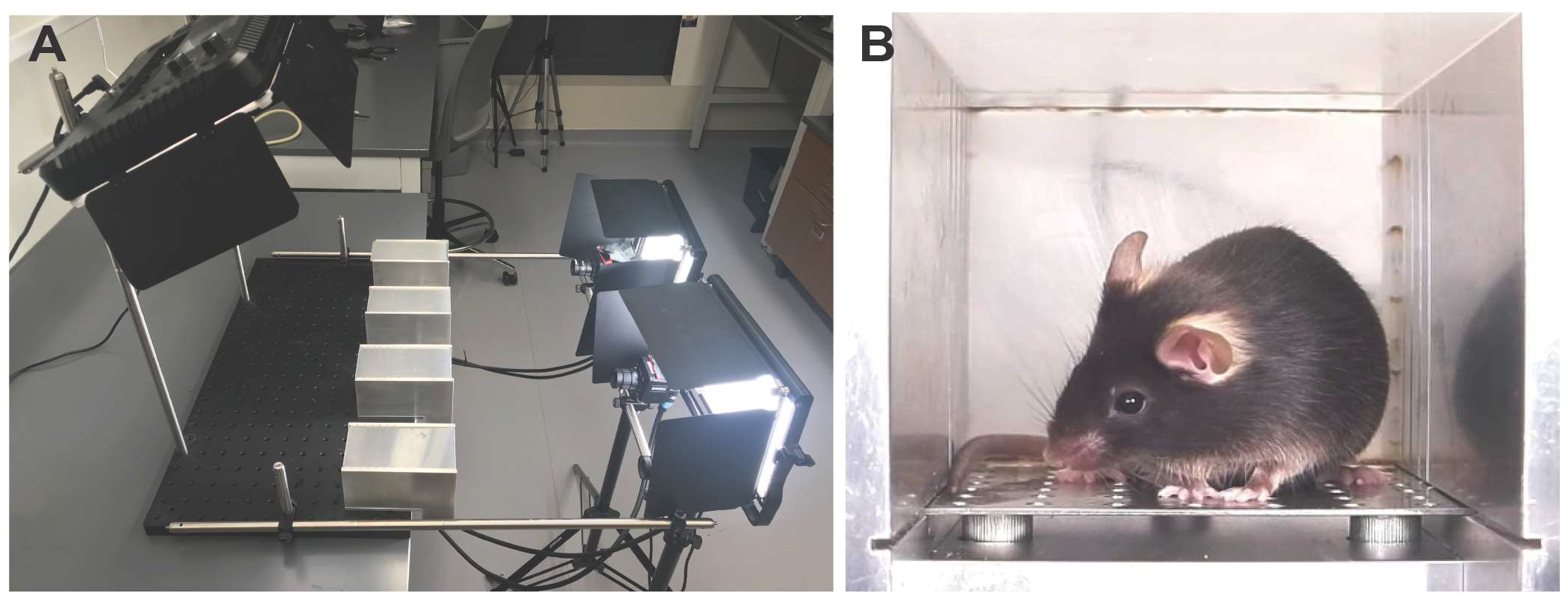
Optimized imaging conditions for black coated mice. (A) Setup showing position of chambers, key lights, back lights, and cameras. (B) Image of properly lit black mouse in imaging chamber. The front is open to the air. Imaging through glass or plastic should be avoided as this creates glare and reflections. Reflections of the mouse can confuse the MLA.

### Black Mouse Grimace Scale

A large amount of human-scored ground truth data is needed to adequately train a machine learning algorithm (MLA). To facilitate standardized scoring by humans, we next developed a black MGS (Figure 2) that is modeled after the original white MGS (Langford *et al*., 2010) and that incorporates observations we made while acquiring >70,000 images of black coated mice (Supplementary Figure 1, Table 1). Three of the facial actions units (orbital tightening, ear position, and whisker change) were scored on a three point scale of “0” (not present), “1” (moderate), or “2” (obvious), after Langford et al. (Langford *et al*., 2010). Nose bulge and cheek bulge action units were difficult to reliably detect and score on this three-point scale, consistent with observations made by others (Hohlbaum et al., 2020). We could confidently determine if a nose bulge was absent (“0”) or obviously present (“2”), allowing us to score nose bulge on a binary scale. It was not possible to reliably detect a cheek bulge, consistent with Hohlbaum et al. (Hohlbaum et al., 2020), so we excluded this action unit from the scale. To calculate the MGS for a single black mouse image, the scores assigned to the four action units are summed, giving a score ranging from 0–8. Scores for individual facial action units can be assigned by expert humans or as described below, by the MLA that we developed.

**Figure 2.**
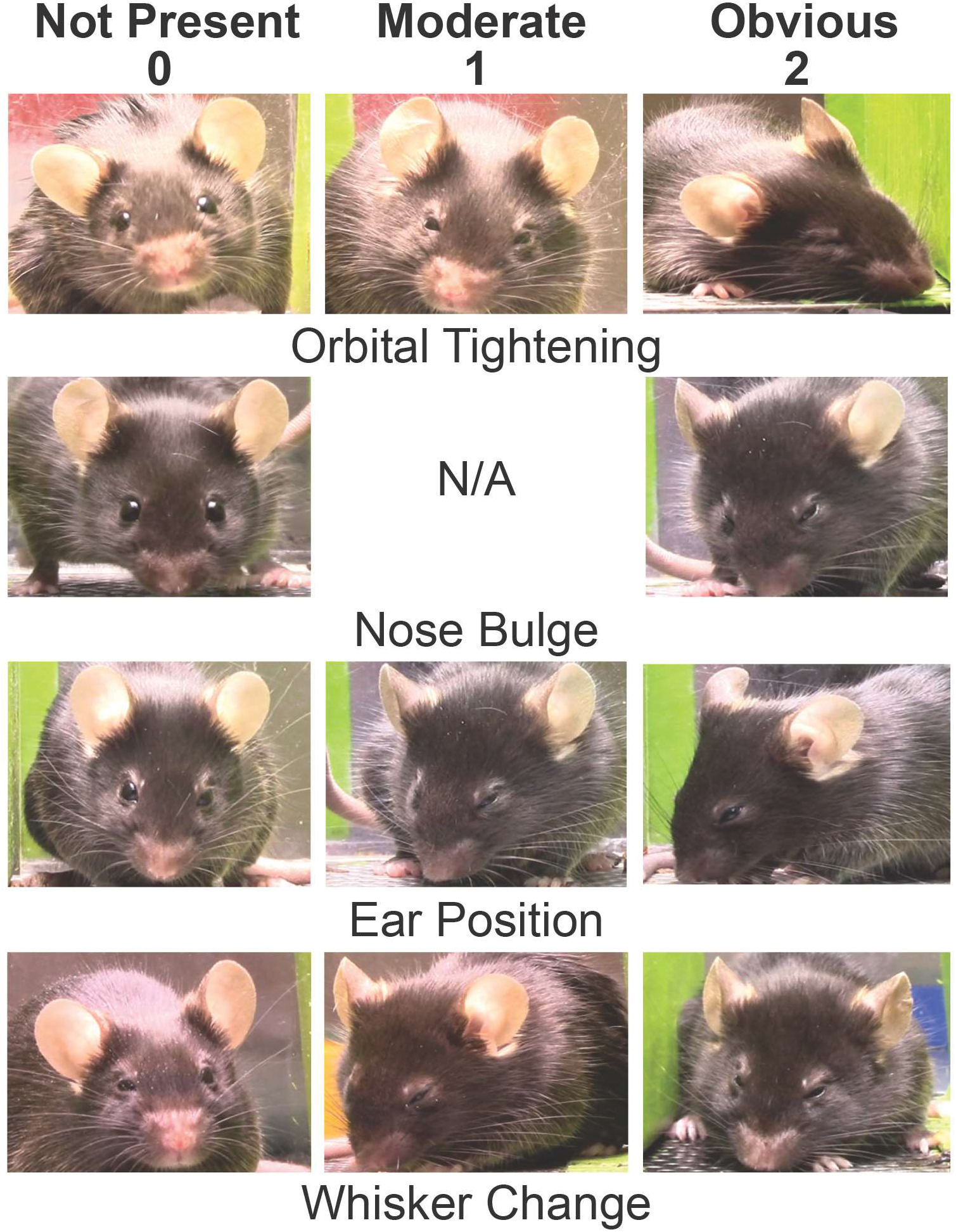
Black mouse grimace scale. The MGS score is calculated for each image by summing the scores assigned to each action unit (0-2 for orbital tightening, ear position, whisker change; 0/2 for nose bulge). Action unit scores can be assigned by humans or by the MLA. N/A, not applicable.

**Table 1.** Number of ground truth images used to train, test, and evaluate the precision of the feature detection model.

|  | **Annotations (train)** | **Annotations (test)** | **Precision** |
| --- | --- | --- | --- |
| **Body** | 72,913 | 9,479 | 99.94% |
| **Face** | 61,380 | 7,978 | 99.63% |
| **Orbitals** | 89,241 | 11,617 | 83.97% |
| **Nose** | 56,591 | 7,393 | 98.47% |
| **Ears** | 136,328 | 17,461 | 99.61% |
| **Whiskers** | 61,209 | 8,238 | 90.59% |
| **Mean average precision** |  |  | 90.19% |

- Precision was computed with Intersection over union (IOU) = 0.5. IOU measures the overlap between the predicted bounding box and the human drawn bounding box. - There are 477,662 human-annotated samples in this data set.

### Generation of the ground truth image dataset

We collected >270 high-definition (1080p, 30 fps, 30 min duration) color videos of C57BL/6 mice under a variety of conditions and states (Supplementary Figure 1), including post laparotomy surgery, a manipulation that reproducibly causes mice to grimace (Langford et al., 2010; Tuttle et al., 2018). We generated a diverse training set to prevent the MLA from overfitting to a specific imaging condition. An array of backgrounds was used, including a background with round holes that looked like eyes (Supplementary Figure 1C,D). Some images contained bedding on the floor (Supplementary Figure 1E) or feces (Supplementary Figure 1F). The color of the side walls was altered to incorporate reflective surfaces and different colors in the training set (Supplementary Figure 1G,H). Videos of mice with ear tags and ear notches, markings that are commonly used to identify animals, were included as well (Supplementary Figure 1I). We also included suboptimal off-center images in the data set (Supplementary Figure 1F,J).

These videos were uploaded to the PainFace software platform (described below), which can extract a user-defined number of random images from a video and has functions to simplify manual (i.e., performed by humans) annotation (drawing bounding boxes around the eyes, nose, ears, whiskers, body, and face) and manual scoring (assigning a score to each action unit, based on the black MGS). We extracted 400–500 random frames per video and manually annotated and scored each image. Many people were needed to annotate and score this large collection of images in a timely manner. To ensure consistency of the resulting ground truth data, annotations and scores associated with each video were spot checked and then electronically “confirmed” (a function in PainFace) by one expert before all the annotations and scores for a given video were added to the ground truth dataset. In total, >475,000 images were annotated and >296,000 images were scored (Tables 1 and 2). This ground truth dataset (MLA build 2021.07.27) was used to train and test the MLA.

**Table 2.** Number of ground truth facial action unit images used for testing and training.

| **Score** | **Orbital** | **Nose** | **Ears** | **Whiskers** |
| --- | --- | --- | --- | --- |
| **No score** | 15,950 | 23,759 | 15,084 | 38,770 |
| **0** | 32,676 | 33,479 | 23,395 | 19,064 |
| **1** | 15,638 | N/A | 27,003 | 13,383 |
| **2** | 9,873 | 16,899 | 8,655 | 2,920 |

### MLA architecture overview

As shown in Figure 3, the MLA consists of two machine learning components: (1) object detection, to locate the face/body from the complex background (Figure 3A) and (2) facial action unit score prediction, which extracts image features around action units and predicts the grimace score (Figure 3B). For the object detection component, we incorporated RetinaNet (Lin et al., 2020), where the input is a video frame and the output includes the top-left and bottom-right coordinates of a bounding box and the category of the object in the bounding box (e.g., face/body; Figure 3A). We used a ResNet50 backbone to identify the object from the input image by hierarchically extracting feature representations in a layer-by-layer manner (He, 2016). We then trained a collection of convolutional neural networks (CNNs) on top of the identified bounding box to assign grimace scores to each action unit (one model per FAU; Figure 3B). Each CNN model outputs confidence values ranging from 0 (low confidence) to 1 (high confidence). The object detection model and grimace scoring models were trained one after another. The flexibility of this modular design makes it possible to add or modify deep models of action units for different mouse strains or for other mammalian species (Dalla Costa et al., 2016; Di Giminiani et al., 2016; Gleerup et al., 2015; Guesgen et al., 2016; Häger et al., 2017; Holden et al., 2014; Keating et al., 2012; Matsumiya et al., 2012; Mogil et al., 2020; Reijgwart et al., 2017; Sotocinal et al., 2011; van Loon and Van Dierendonck, 2015). Similar to other successful machine learning applications such as ImageNet (Deng, 2009), our MLA architecture can accommodate additional action units, such as cheeks which can be scored with white mice.

**Figure 3.**
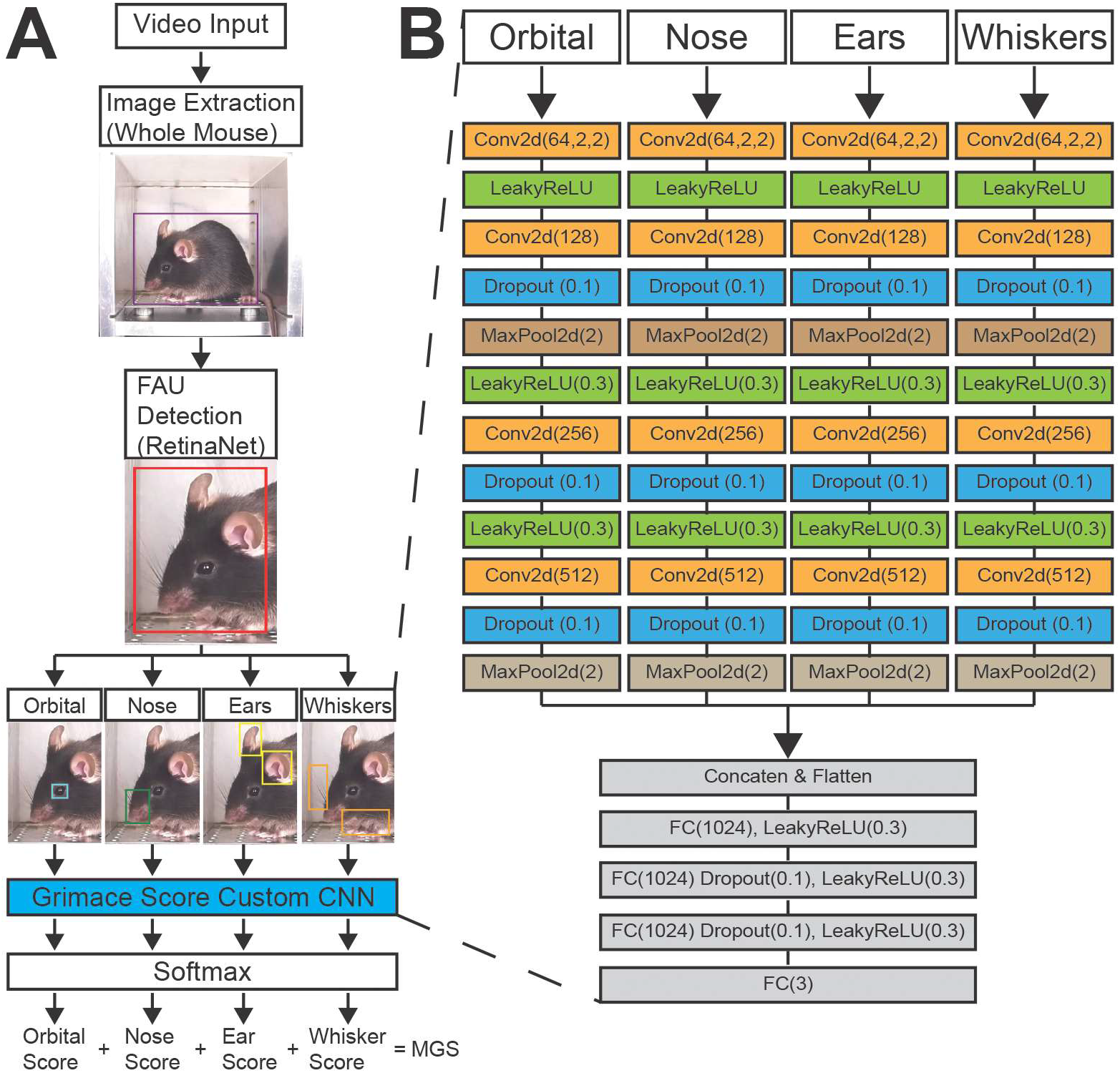
MLA architecture used for training and grimace evaluation. (A) Images are initially processed by RetinaNet with Resnet 50 backbone, which has 50 neuron layers, to detect and annotate the body, face, and each facial action unit (orbitals, nose, ears, and whiskers). (B) Grimace score custom convolution neural network is expanded from (A, indicated by dashed lines). After detection of the body, face, and facial action units, the resulting pixels are processed in a scoring network made up of 18 neuron layers, while each convolutional network has 3 parallel channels with 12 neuron layers. Each facial action unit is processed individually using the scoring network. This process occurs separately for each facial action unit.

### Training and testing the MLA

The ground truth images were separated into training data and testing data, with no overlapping images between these groups (Tables 1-3). For the object detection task, each action unit was treated as a separate class. All possible MGS scores had similar amounts of training data (Table 4) except for the highest MGS score of 8. C57BL/6 mice were rarely assigned a score of 8 under the conditions used in our study.

**Table 3.** Number of ground truth images used to train, test, and evaluate the accuracy of the facial action unit grimace score models.

|  | **Orbital** | **Nose** | **Ears** | **Whiskers** |
| --- | --- | --- | --- | --- |
| **Number of training samples** | 45,764 | 40,168 | 45,184 | 28,180 |
| **Number of test samples** | 11,441 | 10,042 | 11,296 | 7,045 |
| **F-Score** | 0.84 | 0.821 | 0.80 | 0.75 |

- F-score is a composite measurement from the confusion matrix, which considers both sensitivity and specificity of the deep model. It corrects for balancing issues between test and training data sets.

**Table 4.**
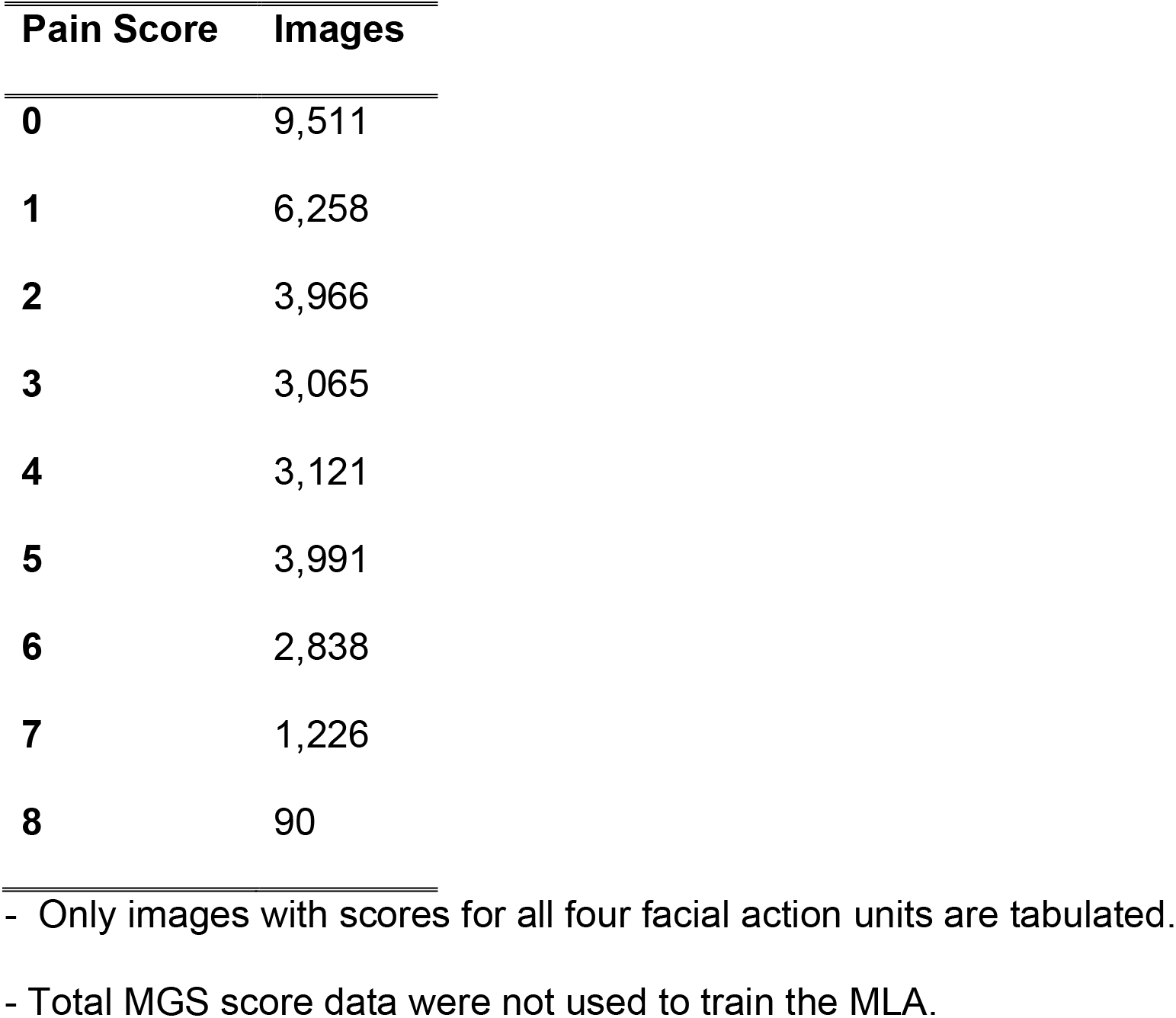
Distribution of human MGS scores in the training data set.

| **Pain Score** | **Images** |
| --- | --- |
| **0** | 9,511 |
| **1** | 6,258 |
| **2** | 3,966 |
| **3** | 3,065 |
| **4** | 3,121 |
| **5** | 3,991 |
| **6** | 2,838 |
| **7** | 1,226 |
| **8** | 90 |

- Only images with scores for all four facial action units are tabulated. - Total MGS score data were not used to train the MLA.

The grimace scoring task was formulated as a classification problem, with each grimace score represented as a class. Mirroring the black MGS (Figure 2), there were three classes for orbitals, ears, and whiskers and two classes for nose. We trained different deep models with similar network architecture to predict the score for each action unit. We did not use a regression approach to estimate each grimace score because this approach tended to revert scores to the global mean. We used three parallel convolutional layers with kernel sizes set to 3x3, 5x5, and 7x7 for feature extraction, and concatenated these networks into a flat vector that projected to a fully connected layer for grimace score classification. The three kernel sizes permit the model to process multiple features from a single image. Although conventional single stream convolutional networks are implicitly agnostic on feature size, our MLA can more clearly separate features, particularly given the different sizes of the FAUs. The negative log likelihood (NLL) function was used for loss estimation (Zhu, 2018). The Adam gradient descent method was used for back-propagation.

After training the face detection model, detection precision with the test images was over 0.99 except for the orbital action unit, which was 0.84 (Table 1). The precision was computed with Intersection over Union (IOU) equal to 0.5. We directly compared the locations of bounding boxes annotated by humans (Figure 4A) to features detected by the MLA (Figure 4B). The MLA consistently identified features in a human-like manner, highlighting the robust feature detection capabilities of the MLA.

**Figure 4.**
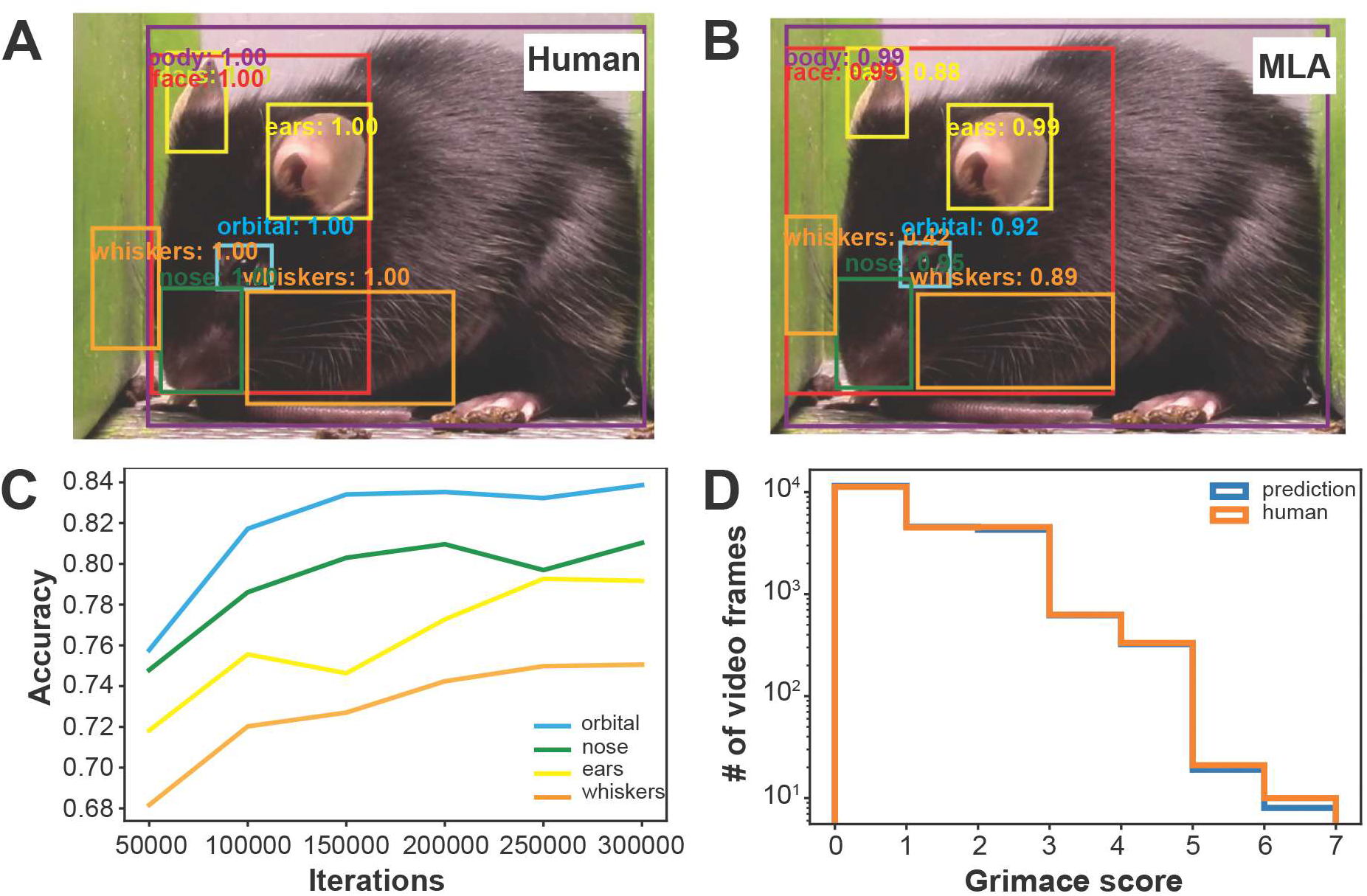
MLA performance. Example showing (A) human and (B) MLA annotation of the same image. (C) Accuracy of the grimace score over MLA training iterations. (D) Human and MLA grimace score concordance.

We also evaluated the accuracy of the MLA at predicting grimace scores. We utilized a confusion matrix to examine the classification imbalance of the grimace scores. The closer to “1” the better the precision of the model. We obtained an F-score of 0.75 or higher for each action unit (Table 3). The accuracy of scoring each action unit improved as the number of training iterations increased (Figure 4C). Moreover, the distribution of grimace scores that were assigned by humans and predicted by the MLA closely overlapped (Figure 4D). The MLA slightly underscored images that humans assigned a high MGS score to, possibly because there were fewer images with a high MGS score in the training set.

We also quantified the difference in human-assigned to MLA-assigned grimace scores on an image-by-image basis (Supplementary Figure 2). The MGS score varied by ± 0.5 for most of the images, with very few images showing a >1 MGS score difference (Supplementary Figure 2A). Similarly, the difference in grimace scores for orbitals, ears, and whiskers action units were in the ± 0.5 range (Supplementary Figure 2B-E). The MLA to human difference was slightly higher for the nose (Supplementary Figure 2C), as expected given that it is much harder for humans to assign a grimace score to the nose of black-coated mice and given the nose is the only action unit that is assigned a binary “0” or “2” score. Overall, these analyses indicate that our MLA accurately identifies where facial action units are located in an image and accurately assigns grimace scores to each facial action unit.

### PainFace software simplifies and standardizes grimace analyses

To make our MLA accessible and easy to use, we created a web-based software platform called PainFace that can be accessed from anywhere in the world with an internet browser (Figure 5A). Users can upload videos (.mp4 format, 30 fps) to a secure server and then initiate a grimace analysis on a GPU workstation. Users can input the “subject type,” “sex,” “data source,” and other metadata that is associated with the experiment. The user can then “initiate grimace analysis” at sampling rates ranging from 0.001 frame/s (low) to 1 frame/s (high). With our current hardware, PainFace can annotate and score up to 300 images per minute, which is >100x faster than an expert human who typically scores ∼3 frames/minute. PainFace can also process multiple videos simultaneously in batches. PainFace generates over two orders of magnitude more MGS data per video than a human, making it possible to analyze grimace data in entirely new ways (see below).

**Figure 5.**
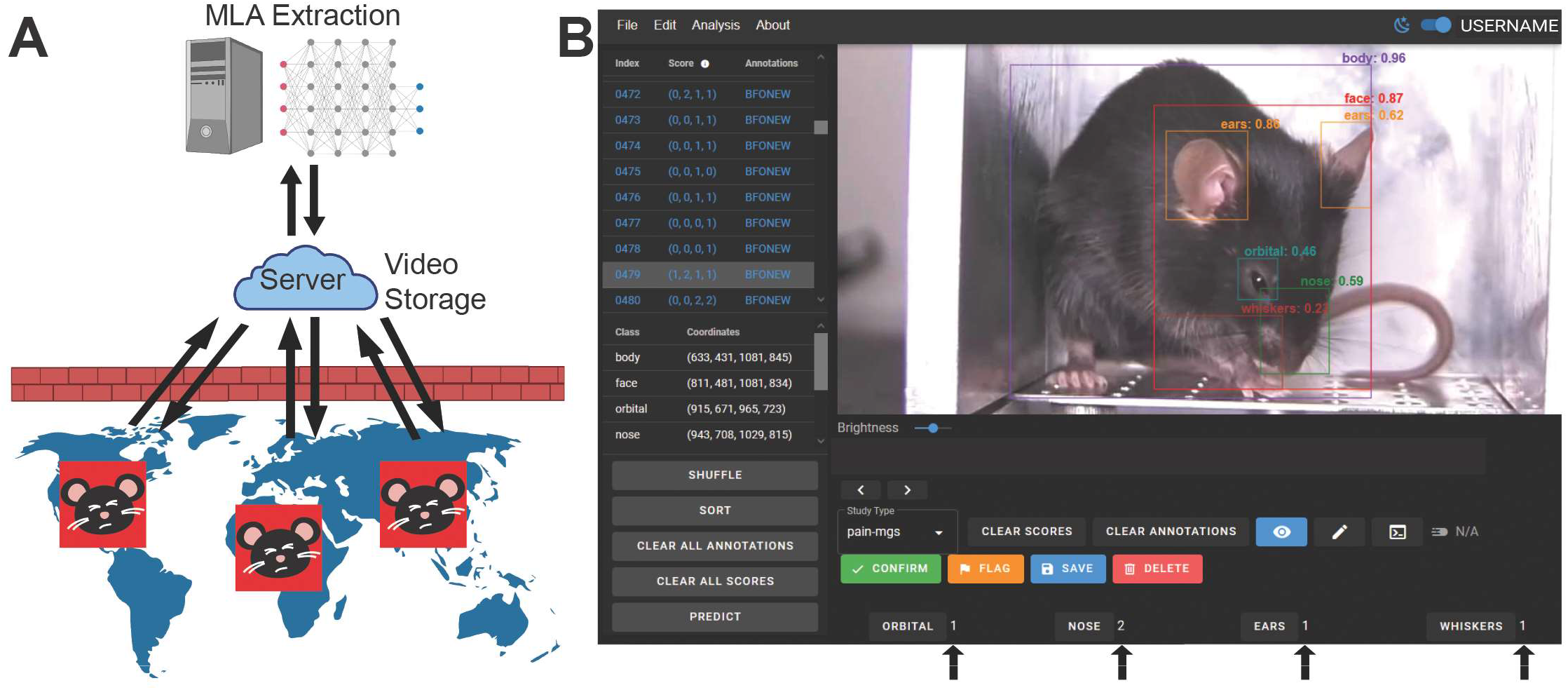
PainFace software platform. (A) PainFace can be accessed from anywhere in the world by visiting http://painface.net with a modern web browser. Videos are uploaded to a secure server for storage. Upon initiating a grimace analysis, the MLA computes a grimace score for every n^th^ (user defined) frame of the video on a GPU cluster. Results are returned to the end-user in a tab-delimited csv file. PainFace users will be given Standard user access (upload videos, initiate grimace analyses, and retrieve analysis results). (B) PainFace screenshot. Bounding boxes and associated confidence values, and scores for each action unit (arrows) are clearly visible.

PainFace generates a comma-separated values (.csv) file containing grimace scores for each facial action unit along with the MGS (sum of each facial action unit score), tabulated for each frame analyzed. The number of facial action units detected per frame is also listed (for black-coated mice, the maximum is four). Additionally, the confidence values for each action unit are listed. The MLA build that was used to complete the analysis is listed in the .csv file. Software build version is important to make note of in publications, as we plan to improve and update the MLA over time. PainFace also generates a .zip file containing the .csv file and the associated image frames that were analyzed. The analyzed dataset can also be opened directly in PainFace, allowing the user to examine the location of the bounding boxes and facial action unit scores frame-by-frame (see example, Figure 5B). While restricted to high-end users, PainFace has tools to annotate and score videos for the purpose of generating ground truth data.

As a cloud-based platform, users can upload videos and perform grimace analyses from any location using an internet browser. Maintaining PainFace in the cloud enhances rigor and reproducibility—everyone has access to the most current software and latest-trained MLA and does not need to install software or worry about software incompatibility issues. Moreover, use of a cloud-based platform supports the goal of creating a standard, easy-to-access platform for grimace analyses. While our current study is focused on black-coated mice, PainFace can be used to generate ground truth data and, once a model is trained, perform grimace analyses with white- or agouti-coated mice or other mammalian species.

### PainFace validation experiment

We next evaluated the extent to which PainFace could be used to quantify grimace following a laparotomy surgery. Prior to surgery, baseline videos were recorded for 30 min from wild-type C57BL/6 male mice. Mice were then separated into three groups (Figure 6A): 1. sham surgery, 2. LAP (laparotomy surgery), 3. CARLAP (the analgesic carprofen was administered just prior to making the incision). The mice were placed on a heated pad for 30 min following surgery. The mice were then returned to the imaging chamber, and videos recorded for 30 min. Baseline and post-sham/surgery videos were uploaded to PainFace.

**Figure 6.**
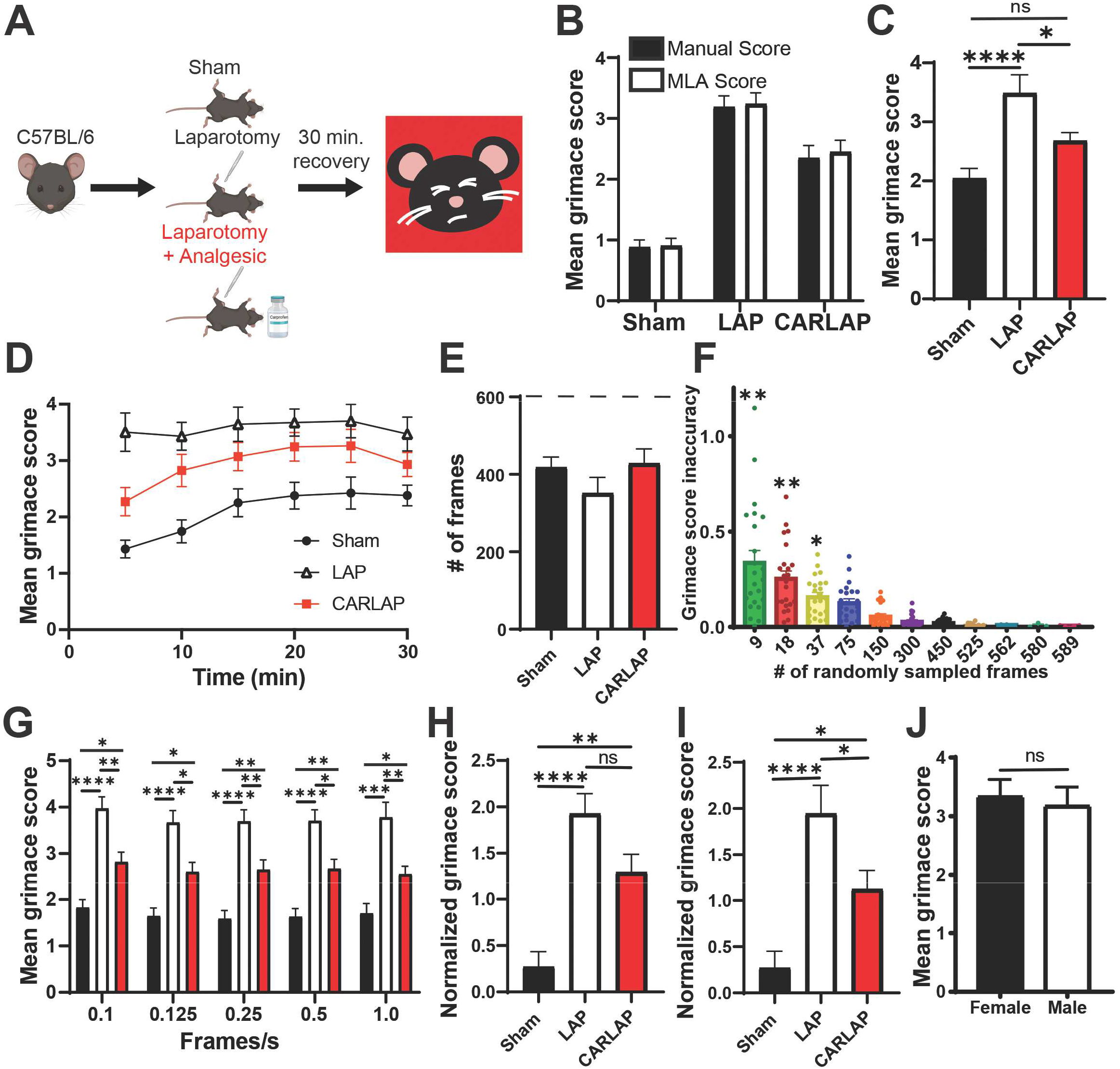
Validation experiment. (A) Videos (30 min. duration, 30 fps) from the sham, LAP, and CARLAP experimental groups were manually scored by humans and by the PainFace MLA. (B) No significant differences between human and MLA-scored videos (within group comparisons; from the whole 30 min. video, ten random frames containing four action units were scored from 12 validation videos per group). (C) Mean PainFace grimace score from each 30 min. video (1 frame evaluated by PainFace from each second of video = 1,800 evaluations per video, only images with all four action units used). (D) Mean PainFace grimace score plotted in 5 minute bins (150 random frames sampled per bin containing all four action units) over the 30 min. video. (E) Mean number of frames (out of 600 frames analyzed, dashed line) where all four action units were detected and scored by PainFace during the first 10 min. (F) Analysis of grimace score inaccuracy relative to the number of frames analyzed during the first 10 min. Calculated by subtracting the mean grimace score of (every frame with four action units scored by PainFace) minus (the indicated # of frames with four action units scored by PainFace); frames collected from the first 10 min. of each video. (G) Mean grimace score after evaluating the first 10 min of each video with PainFace at the indicated frame rates. (H) Mean grimace score from 30 min. videos normalized to baseline videos. (I) Mean grimace score for first 10 min of each video normalized to first 10 min. of baseline video. (J) Mean grimace score of males and females post LAP (n=11 males, n=9 females) from first 10 min. of video. (B-I) n = 12-17 male mice/group. *p < 0.05, **p < 0.01, ***p < 0.001, ****p < 0.0001. Statistics: Brown-Forsythe ANOVA, Kruskal-Wallis, or T-test for male:female comparisons.

For the initial analysis, we sought to compare MLA scores to human scores. Manual scoring is time consuming and labor intensive for humans, so it is customary to randomly select 8-10 frames from each video for manual scoring (Langford et al., 2010; Sotocinal et al., 2011). We thus extracted 10 random frames containing all four action units (i.e., indicates mouse is facing the camera) from each video and scored manually and with PainFace (MLA build 2021.07.27 was used to score these 12 videos from each group for a total of 36 videos). Manual and MLA scores were not significantly different within the experimental groups (Figure 6B), indicating concordance between human and PainFace scores. We next analyzed the 30-min videos at a high 1 frame/s sampling rate with PainFace (Figure 6C). As expected, the mean MGS score of LAP animals was significantly higher than sham surgery animals, and the mean MGS score was attenuated in mice treated with an analgesic just prior to the surgery (CARLAP group). We noticed that grimace scores in the sham group were approximately one unit lower when scored by PainFace at 10 frames/video (Figure 6B) versus when scored at ∼1,800 frames/video (Figure 6C). We speculate that the low sampling rate has the potential to under-sample bouts of grimacing (which would reduce the overall score) relative to the high sampling rate. As another possibility, the low sampling rate may not accurately capture the grimace score from the 30-min videos if grimacing (or lack of grimacing) varies over time.

To better evaluate the dynamics of grimace over time, we plotted the mean MGS score of each experimental group in 5-min interval bins. We found that grimace scores for the LAP mice remained uniformly high for the entire 30 min video (Figure 6D). However, the sham and CARLAP groups were not uniform over time. The largest difference in grimace scores between the sham, LAP, and CARLAP groups occurred within the first 10 min. Since this time window appeared to be optimal for detecting differences in MGS scores post laparotomy, we focused subsequent analyses on this 10-min window.

Within the first 10-min window, PainFace detected 350-450 frames with 4 FAU out of 600 frames analyzed from each group (Figure 6E), suggesting the animals faced the camera ∼65% of the time (in general, PainFace only detects all four action units if the mouse is facing the camera and if the face is well-illuminated). However, there was a range of 4 FAU frames, ranging from 0-598 per animal. We thus sought to determine how many 4 FAU frames must be averaged to calculate an accurate MGS score. We randomly sampled frames from 23 videos with 500 or more 4 FAU frames in the first 10 minutes. We then plotted the MGS score inaccuracy as a function of the number of randomly sampled 4 FAU frames (Figure 6F). Inaccuracy was determined for each video by calculating the difference between the sampled grimace score (for example, the average of 9 or 75 randomly sampled frames) and the video’s true grimace score (i.e. the average grimace score from ≥500 frames from that video). We found that a minimum of 75 frames over the 10 min window (7.5 frames/min = 0.125 frames/s) was sufficient to get an accurate grimace score. For comparison, humans manually score videos at a much lower rate (1 frame with mouse facing forward is randomly identified and scored every 3 minutes; 0.006 frames/s) (Langford et al., 2010; Sotocinal et al., 2011). As our analysis shows (Figure 6F), low sampling rates have the potential to introduce higher variability into manual MGS score estimates when compared with appropriately powered automated grimace analyses using PainFace.

We next analyzed the validation videos at intervals ranging from 0.1 frame/s to 1 frame/s with PainFace (4 FAU frames only), to bracket the sampling rates that produced accurate grimace scores (Figure 6G). The sham versus LAP groups and LAP versus CARLAP groups were significantly different from one another at all sampling intervals (Figure 6G), indicating that sampling at 0.1 frame/s to 1 frame/s is sufficient to generate accurate grimace scores and to detect differences between experimental groups. While PainFace can evaluate videos at any frequency, including every frame of a 30-fps video (three orders of magnitude greater than a human), we opted to simplify the interface by giving users the option to analyze videos at sampling rates of 0.001 frame/s (low, useful for assessing overall video quality and percentage of frames with 4 FAU before sampling at a higher rate), 0.1 frame/s, and 1 frame/s. We may limit or remove the computationally intensive 1 frame/s option in the future if user demand exceeds PainFace computational resources.

We also analyzed the first 10 minutes of these videos using less stringent filtering criteria: 1+ FAU/frame (one or more FAUs detected per frame), 2+ FAU/frame, and 3+ FAU/frame. No significant differences were seen when compared to using the 4 FAU/frame filtering criteria (data not shown). This is likely due to the high proportion of 4 FAU frames in our validation data set (Figure 6E).

We also detected significant differences between the sham versus LAP groups and LAP versus CARLAP groups when MGS scores from the 30-min videos (Figure 6H) and 10-min videos (Figure 6I) were normalized to baseline MGS scores. Lastly, PainFace detected similar levels of grimace in male and female mice post laparotomy (Figure 6J), indicating that PainFace can be used to quantify facial grimaces in black-coated mice of either sex.

### Distribution of grimace scores

PainFace generates a large amount of data relative to manual scoring. To explore these data at a granular level, we plotted the distribution of MGS scores from 6,000 individual frames from each experimental group (Figure 7A). We found that these data were not normally distributed (Supplementary Figure 3A-D). The frequency distribution was skewed left towards lower MGS scores in the sham group and skewed right towards higher MGS scores in LAP animals (Figure 7A, Supplementary Figure 3A,B). The CARLAP frequency distribution was skewed slightly left toward lower MGS scores (Figure 7A, Supplementary Figure 3C), further indicating that carprofen reduces grimacing following surgery (Figure 7A). Statistical differences were also detected in the magnitude of skew between the sham and LAP groups (Supplementary Figure 3E). Comparison of kurtosis values across groups did not yield statistical differences (Supplementary Figure 3F).

**Figure 7.**
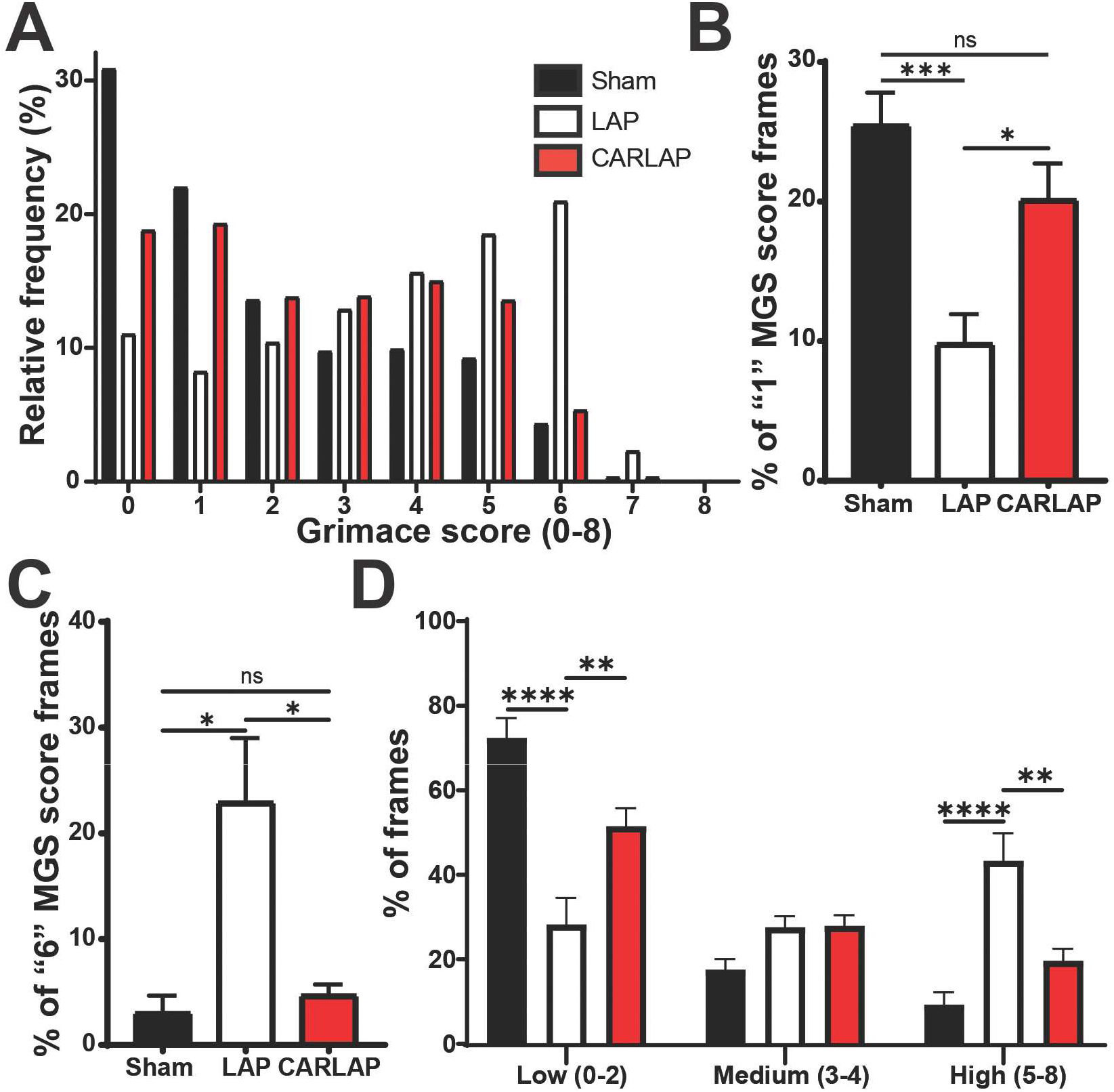
Histogram analyses of validation data. (A) Distribution of PainFace grimace scores (0-8) across all 30 min. of the videos (10,000 frames with 4 FAU/group). (B) Percent of frames with a grimace score of 1 (across first 10 minutes of each video). (C) Percent of frames with a grimace score of 6 (across first 10 minutes of each video; 6,000 frames with 4 FAU/group). (D) Distribution of MGS scores in low, medium, and high categories (sampled from first 10 minutes of each video). n = 14-17 male mice/group. *p < 0.05, **p < 0.01, ***p < 0.001, ****p < 0.0001. Statistics: Brown-Forsythe ANOVA test.

We next plotted the percentage of frames with low (“1”) MGS scores and high (“6”) MGS scores for each experimental group (Figure 7B,C). This analysis likewise indicated that LAP animals spent significantly less time in a low grimace state and ∼7x more time in a high grimace state relative to sham and CARLAP animals (Figure 7B,C). We also binned frames into low (0-2), medium (3-4), and high (5-8) MGS score categories (Figure 7D). Approximately 75% of frames from the sham group fell into the low grimace category, with only ∼10% falling in the high grimace category (Figure 7D). In contrast, 45% of the frames from LAP mice fell within the high grimace category. CARLAP animals were skewed towards the low grimace category (Figure 7D). Collectively, these data indicate that animals spend more time in a high grimace state after undergoing a surgical procedure and an analgesic drug can reduce the amount of time animals spend in this high grimace state.

### Specific facial action score combinations are overrepresented in mice that grimace

Grimace scores can be decomposed to unique combinations of individual FAU scores. Thus, we next quantified the relative frequency of each FAU combination in mice from each experimental condition (Figure 8). The most frequent score combinations in the sham group were 0-0-0-0 (orbital-ears-whiskers-nose), 0-0-1-0, and 0-1-0-0. Specific FAU combinations (1-1-1-2 and 2-1-1-2) were upregulated in LAP group relative to sham controls and reduced in the CARLAP group relative to the LAP group. These data suggest that specific grimace score combinations may serve as a facial signature or expression of pain.

**Figure 8.**
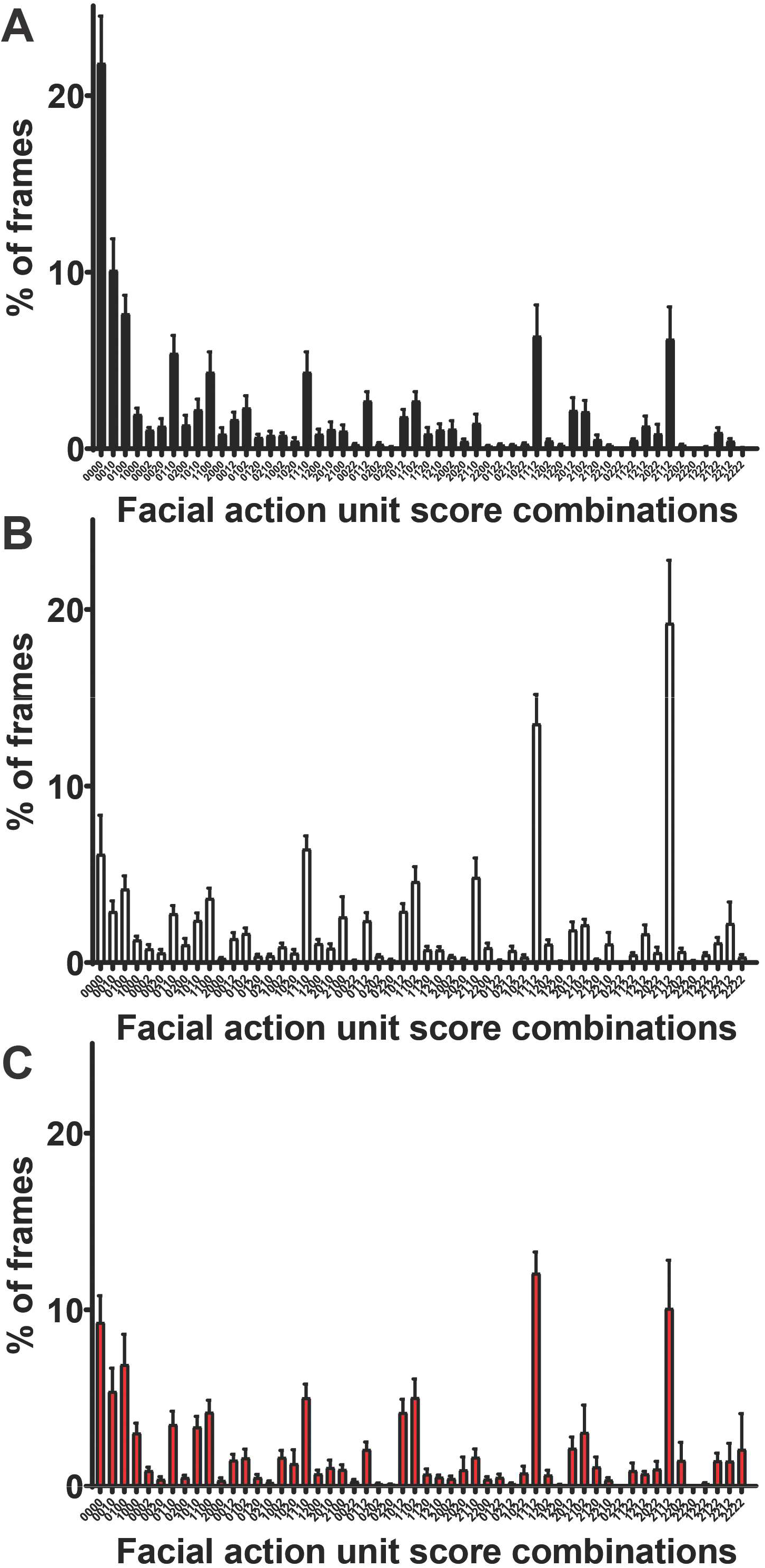
Frequency distribution of facial action unit score combinations for each frame of (A) sham, (B) LAP, and (C) CARLAP experimental groups. OEWN = orbital, ears, whiskers, and nose facial action units.

### PainFace replication experiment in the Mogil lab

We collaborated with Dr. Jeffrey Mogil at McGill University in Montreal, Canada to determine if PainFace can be used by another lab to quantify facial grimaces. The Mogil lab found that the inflammatory agent carrageenan causes mice to grimace when bilaterally injected into both hind paws (Langford et al., 2010). To determine if the imaging setup and/or light levels (∼1,500 lx) artificially reduced grimace, possibly via stress-induced analgesia (Butler and Finn, 2009), one group of mice was habituated to the imaging setup and lighting conditions for 1 h each day for 3 days prior to carrageenan injection, while another group was not habituated (Figure 9A). On the day of the experiment, the mice were placed in the imaging setup for 30 min and then a 30-min baseline video was recorded. The mice were then injected with 1% carrageenan (bilaterally, both hind paws) and a 1-h video was collected from each mouse 3 h post-injection. These videos were manually scored by the Mogil lab (10 frames scored/30 min video) and by PainFace (1 frame/s). Both scoring approaches (manual and PainFace) detected a significant increase in the MGS score from 3.5-4 h post carrageenan injection relative to baseline in the not-habituated animals (Figure 9B). PainFace detected a significant increase in the MGS score during this same time period in the habituated animals (Figure 9B). No significant differences were observed between BL and 3.0-3.5 h post carrageenan in habituated or not-habituated animals using manual or MLA scoring (data not shown). Overall, these experiments demonstrate that MGS scores generated by PainFace are in agreement with scores generated manually by the Mogil lab; which is located at a different university, in a different country, with different housing conditions, building environments, and experimenters.

**Figure 9.**
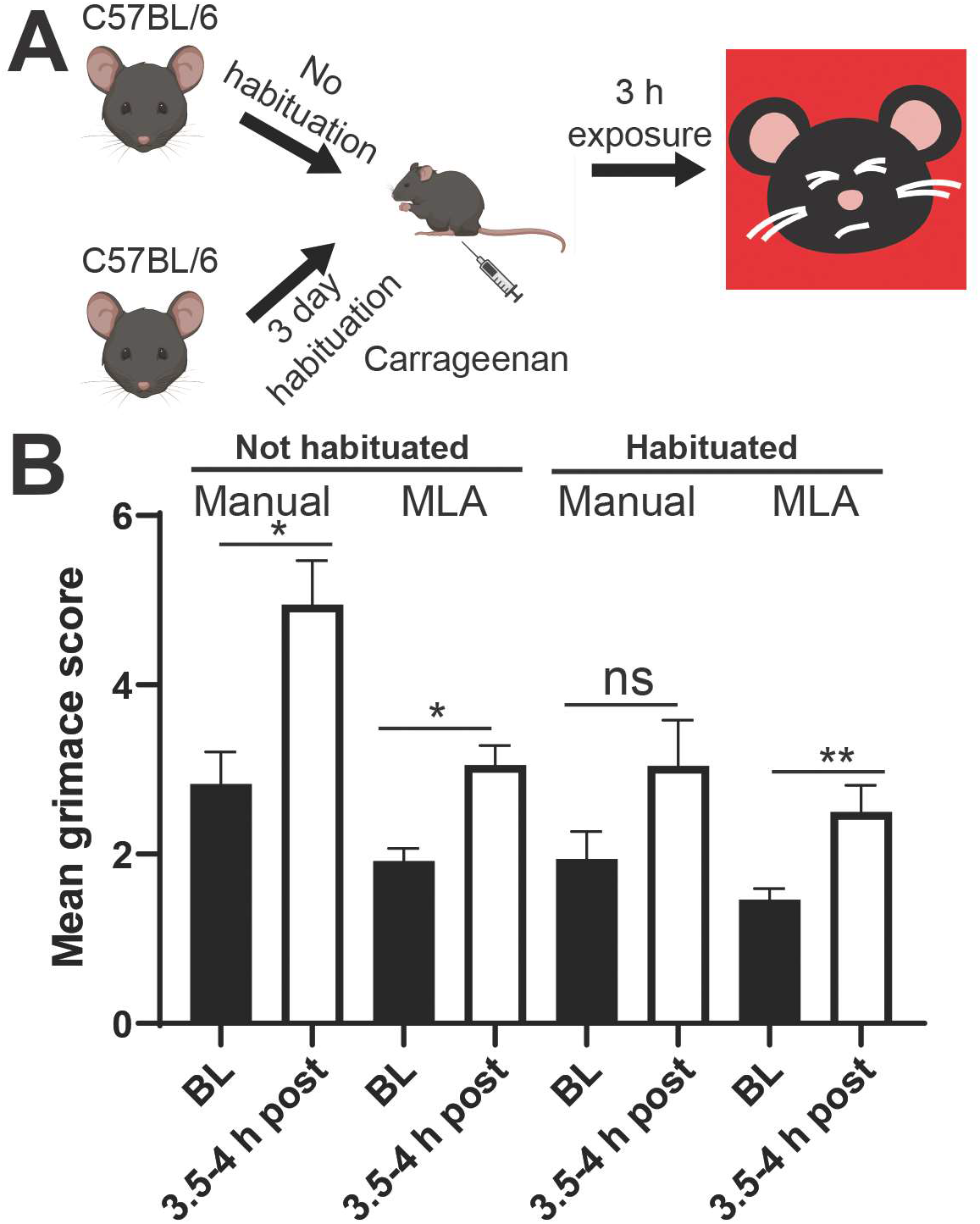
Mogil lab validation experiment. (A) Mice were habituated or not habituated to the setup and lighting, and then were injected with 1% carrageenan (bilateral, both hindpaws) in Dr. Jeff Mogil’s lab at McGill University, Canada. (B) Videos (1 h) were collected 3 h post-injection and scored manually by the Mogil lab and scored with PainFace (MLA). n=10 male C57BL/6 mice. Statistics: Brown-Forsythe ANOVA.

### Inter-rater reliability of humans relative to PainFace

Rodent grimace scales feature high inter-rater reliability within individual laboratories (Langford et al., 2010). Over the course of this research, the Zylka and Mogil labs became proficient in scoring images with the black-coated MGS (Figure 2). To determine the reliability of this scale, a randomized set of 25 images was generated by the Mogil lab and these images were then scored by four expert scorers across the two labs (Zylka lab, n=3; Mogil lab, n=1;). We found general agreement in the total grimace score, evaluated by the intraclass correlation coefficient (ICC), across the Zylka lab (ICC_Average-ZylkaLab_ = 0.939; Table 5) and the two laboratories (ICC_Average-BothLabs_= 0.765; Table 6). The ICC_Average-ZylkaLab_ was similar to the previously reported intra-lab reliability in grimace scoring (Langford et al., 2010).

**Table 5.** Inter-rater reliability for Zylka lab.

| **Feature** | **ICC Single Measure** | **ICC Average Measure** |
| --- | --- | --- |
| Orbital | 0.76 | 0.91 |
| Nose | 0.61 | 0.83 |
| Ears | 0.93 | 0.97 |
| Whiskers | 0.36 | 0.63 |
| Total Score | 0.84 | 0.94 |

**Table 6.**
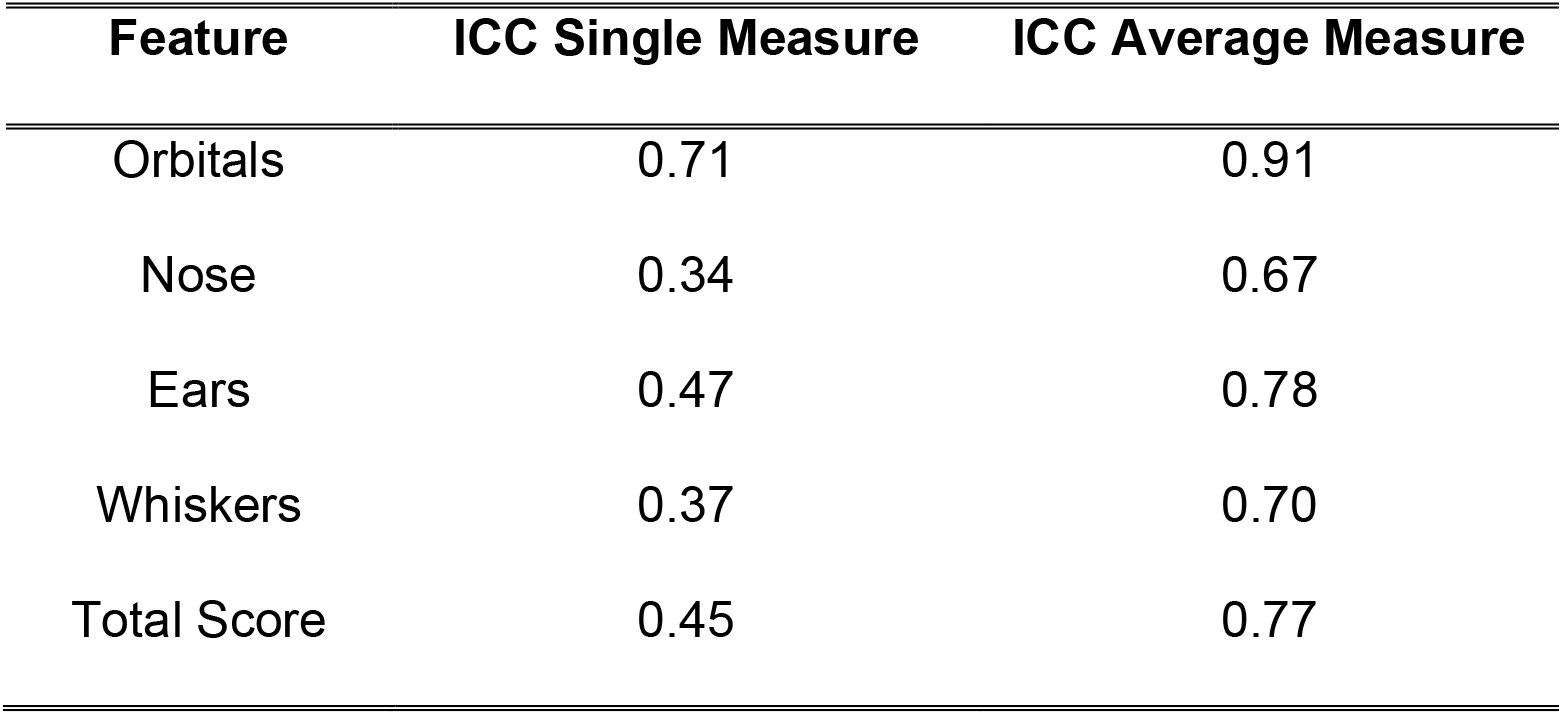
Inter-rater reliability for Zylka and Mogil labs.

| **Feature** | **ICC Single Measure** | **ICC Average Measure** |
| --- | --- | --- |
| Orbitals | 0.71 | 0.91 |
| Nose | 0.34 | 0.67 |
| Ears | 0.47 | 0.78 |
| Whiskers | 0.37 | 0.70 |
| Total Score | 0.45 | 0.77 |

We also examined the ICC of individual FAUs both at intra- and inter-laboratory levels. Scoring reliability was not uniform for each FAU (ICC_Average-ZylkaLab_ range: 0.63 – 0.97 and ICC_Average-BothLabs_ range: 0.67 – 0.91). The most unreliable FAU scores across the two laboratories were the nose and whiskers. Inter-lab variability when scoring individual FAUs has the potential to negatively impact reproducibility across labs. In contrast, PainFace provides a way to standardize grimace scoring across laboratories, regardless of location or level of scoring proficiency.

## Discussion

Manual scoring of facial grimaces requires a significant investment in training personnel, many of whom leave the lab creating a need to train more people, and it is challenging to collect high-quality videos of the mouse face, especially from black-coated mice. The reliance on human experts, who may have different levels of proficiency and/or styles in scoring (Table 5 and Table 6), can be perceived as subjective and less quantitative when compared to evoked assays that measure withdrawal responses in units of time, force, or frequency.

In a prior attempt to reduce subjectivity when scoring facial grimaces, we developed a MLA that outputs a binary pain/no-pain assessment with images of white-coated mice (Tuttle et al., 2018). However, the algorithm was not designed to output numerical MGS scores nor did the algorithm work with black-coated C57BL/6 mice, the predominant strain used by preclinical pain researchers (Sadler et al., 2022). We made the software available on Github (
https://github.com/BenjaminCorvera/MGS-pipeline-distribution), along with detailed instructions on how to install and use the software. However, the software must be operated through the command line interface, which is not intuitive for most neuroscientists, creating a barrier to wider use. Subsequently, other groups developed automated ways to quantify grimace responses in white or black-coated mice (Andresen et al., 2020; Chiang et al., 2022; Rea et al., 2022). However, image diversity was low, which is known to cause machine learning models to overfit to the specific imaging conditions used. Moreover, no user-friendly software or hardware has been developed, further limiting broad utility.

Initially, we found that it was difficult to collect high-quality videos of black-coated mice using the lighting conditions that worked well for white mice (Tuttle et al., 2018). To overcome this barrier, we developed an inexpensive imaging setup made from commercially available parts (Figure 1). We strongly encourage labs to use this setup, lighting, and one of the recommended cameras when acquiring videos of black-coated mice. Imaging through air eliminates glare that commonly occurs when imaging through glass or plastic.

To overcome the barrier of subjectivity in scoring, we generated a large ground truth data set and used these data to train convolutional neural networks to detect and score each facial action unit. The MLA outputs numerical grimace scores that are on par with expert scorers (Table 3). To eliminate the need to use the command line interface, we developed a user-friendly software platform called PainFace. This cloud-based software can be accessed from anywhere in the world with a modern web browser, and all commands are easy to execute via a graphical user interface. These innovations simplify and standardize grimace scoring across laboratories, as demonstrated by replication experiments performed in two labs located in different countries. Individuals within labs and across labs do not consistently score the same images the same way (Tables 5 and 6), highlighting the need for a standardized platform to enhance rigor and reproducibility in grimace analyses and pain research more generally.

As with any research tool, there are limitations to keep in mind when using the PainFace software platform. First and foremost is video quality. PainFace will output MGS scores even if images are poor quality (such scores may be associated with low confidence values). Things that can degrade MLA performance include low light levels, unfocussed video, mouse not facing the camera, reflections of the mouse face, additional mice in the video, and round dark objects that look like eyes. As a rule of thumb, if any of the facial action units are hard for a human to see, these action units will be hard for the MLA to detect as well. Digital cameras can also vary in terms of their optics and image sensors, which can impact image characteristics that are imperceptible to humans but that can nonetheless confuse MLAs. It is thus important to use a camera and video resolution that is similar to what was used to generate the ground truth data (see Methods). As noted above, the current MLA build tends to underscore relative to humans. From a practical perspective, this means that PainFace may not detect grimacing if a manipulation causes low levels of grimacing, as assessed by humans. We performed laparotomy surgeries and carrageenan injections because these manipulations produce high grimace scores (Langford et al., 2010). Machine-learning models tend to improve as more training data is generated (Shankar et al., 2020), so there is the potential that underscoring can be resolved with more training data.

We spent considerable effort adjusting lighting so that all facial features of black-coated mice were accurately captured. Despite these efforts, we recognize that this variable could be optimized further. Light intensity, wavelength (“temperature”), and flicker frequency can differ depending on the light sources used. Cool (4000K) light color and >60 lx light intensity reportedly increased anxiety in mice (Castelhano-Carlos and Baumans, 2009; Kapogiannatoua, 2016). For comparison, light intensity in labs is typically 750-1200 lx (https://www.archtoolbox.com/recommended-lighting-levels/). We did not control these variables when collecting the ground truth data, as we empirically found that mice grimaced under the lighting conditions that we used. Subsequently, we tested additional lighting conditions using a top light and key light and measured the light intensity within the chamber (∼1,500 lx). Mice grimaced under these conditions and videos from these mice were scored with PainFace (see Figure 9). We do not recommend recording videos of black-coated mice under very low light intensities and then increasing brightness with video editing software as this manipulation has the potential to compromise PainFace feature detection and scoring accuracy.

We recognize that investigators will want to use PainFace in other settings and with other mouse strains. To facilitate improvements to the MLA over time, we plan to identify user-uploaded videos (Figure 5) that differ in some way from the videos in the current ground truth data set. We will annotate and score images from these videos and then incorporate them into the ground truth data set. For example, some users may opt to collect videos in a different type of imaging chamber, which we do not recommend, or collect videos of mice with headcaps, conditions that we did not use. We intend to re-train the MLA as more ground truth data is acquired. Thus, investigators should note which MLA build was used and avoid inadvertently using two different builds when analyzing data from a single study or experiment. We recognize that PainFace has the potential to be used by many labs for grimace analyses. We urge researchers who use PainFace to not become complacent, and to occasionally check the output of PainFace relative to manual scoring and relative to negative (baseline or sham) and positive controls (such as LAP animals). If PainFace output does not meet expectations, video frames can be flagged (a function in PainFace), allowing us to inspect the flagged frames to determine what might have caused the misclassification.

PainFace can score substantially more images from videos than humans, unlocking new ways to analyze grimace data. As a case in point, we analyzed the distribution of grimace scores from 6,000 frames, collected from the first 10 min of three different experimental groups. This data-driven analysis led to the identification of low, medium, and high grimace states, and the observation that sham, LAP, and CARLAP animals spent different amounts of time in these states (Figure 7). CARLAP animals distributed more towards the low and medium grimace states relative to the LAP animals. This type of analysis suggests that carprofen, and perhaps other analgesics, may not simply blunt nociceptive signals or “reduce pain.” Instead, analgesics may facilitate transitions to other internal states, or stabilize other internal states, akin to how attending to a non-painful stimulus reduces the perception of pain (Bohic et al., 2021; Hodes et al., 1990; Miron et al., 1989; Siedenberg and Treede, 1996). We also found that unique combinations of facial action unit scores were overrepresented when mice were in a high grimace state—an observation that could not readily be made with a small number of human-scored images. Unique facial action unit score combinations may be indicative of specific behavioral states, analogous to how different facial expressions denote specific emotional states in humans and mice (Dolensek et al., 2020; Horstmann, 2003; Smith et al., 2018).

### PainFace beta testers, a new approach to enhance rigor and reproducibility

To further enhance rigor and reproducibility, we will make PainFace software available to a small number of experts in the pain field, to beta test the imaging setup and the software prior to public release. We will solicit beta testers via social media and reach out to investigators that requested a MLA for black mice after reading our white-coated mouse MLA study (Tuttle et al., 2018). We will offer training sessions on how to collect images and on how to use the software, and we will provide each beta tester with one PainFace user account (standard user permission level). Beta testers will have the opportunity to use the software for grimace analyses. We will ask that beta testers report bugs and provide suggested improvements to the user interface.

We may use videos uploaded by beta testers to improve the MLA and increase the size and diversity of the training data. For example, none of our training data includes images of mice with head caps or cables/wires attached to their head. We currently do not know if PainFace will accurately identify facial action units when mice have things attached to their head. Should PainFace have difficulties with such images, these images can be flagged. One of our experts can then randomly select hundreds of images from the flagged videos and manually generate training data for inclusion into the ground truth data set. Following the beta period, we will re-train the MLA using additional ground truth data sourced from our lab and from beta testers, update the MLA build, revise the tables presented above, and submit the manuscript for peer-review to an open access journal. We plan to acknowledge beta testers for their contribution if they give us permission to do so. By enlisting experts in the pain research field as beta testers, this will help us to identify issues that could impact rigor, reproducibility, and/or platform usability prior to peer review and before wide release.

## Acknowledgements

This work was supported by a grant from the NINDS (R01NS114259) to M.J.Z. NSF GRFP awarded to R.P.P.

## Contributions

ESM, SKP, RPP, GW, and MJZ participated in the study conceptualization/design and wrote the paper. ESM, DFR, JEL, BTB, JLK, XH, RLG, ADK, JAJ, SH, AKT, KAV, RMM, HOB, and MRK provided ground truth grimace annotations and/or scores for MLA training. Experimental work in the Zylka lab was conducted by ESM, RPP, and JEP. Optimized hardware and software for data acquisition were developed and implemented by RPP. SKP and GW developed the PainFace MLA and server platform. ZJM and JJN developed the web-based user interface application. All data analysis was conducted by RPP and overseen by GW and SKP. Replication experiments were overseen by JSM and conducted by SGS, LVL, and J-SA. MJZ and JSM oversaw inter-lab reliability experiments conducted by ESM, DFR, RPP, and SGS.

## Supplementary Figures

**Supplementary Figure 1.**
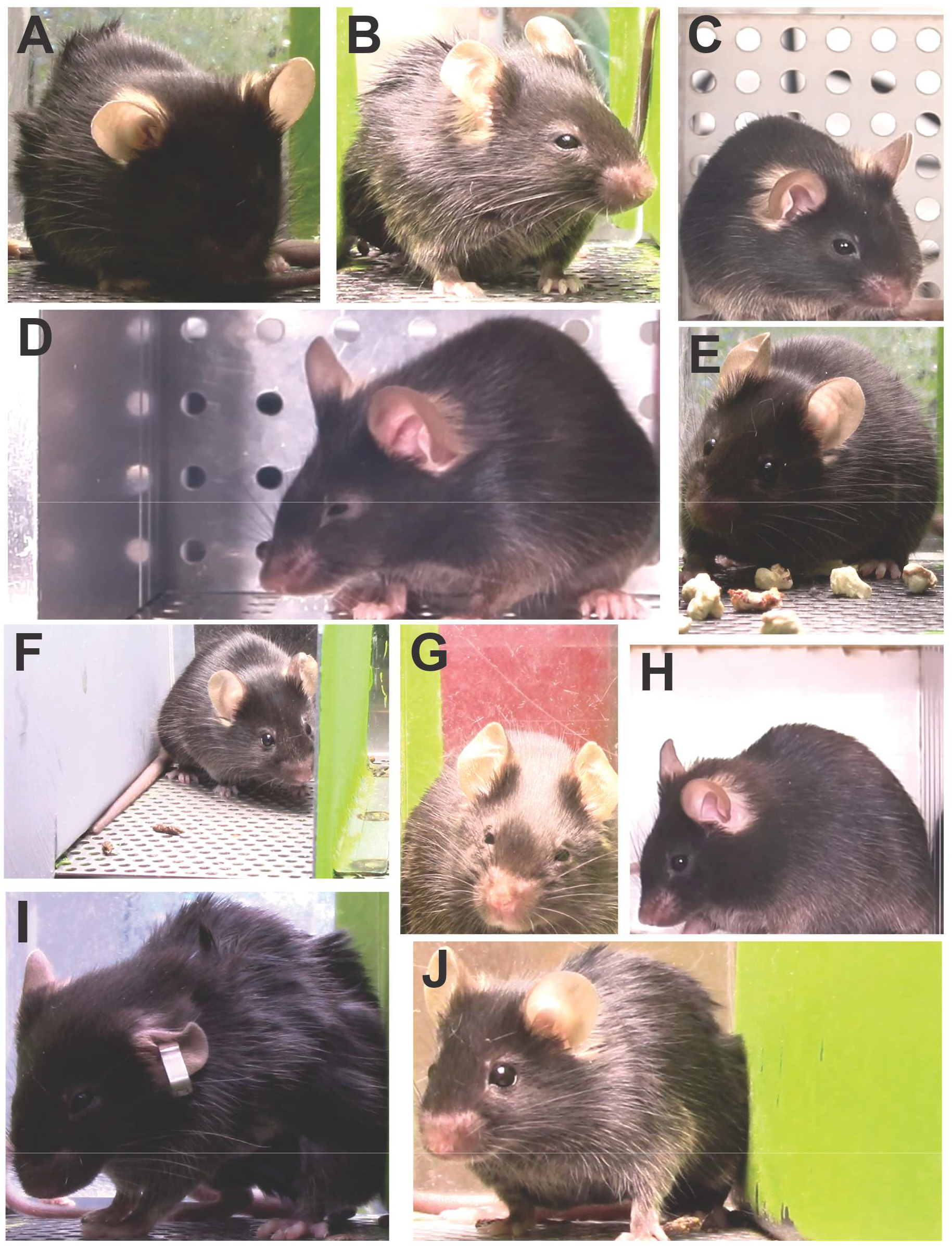
Example images in the training set. (A) Too dark, (B) clear, well-lit, (C,D) metal background with holes that an early build of the MLA misclassified as eyes, (E) bedding on floor, (F) angled camera with gray wall and feces on floor, (G) red background, (H) opaque background, (I) ear tags, and (J) off center.

**Supplementary Figure 2.**
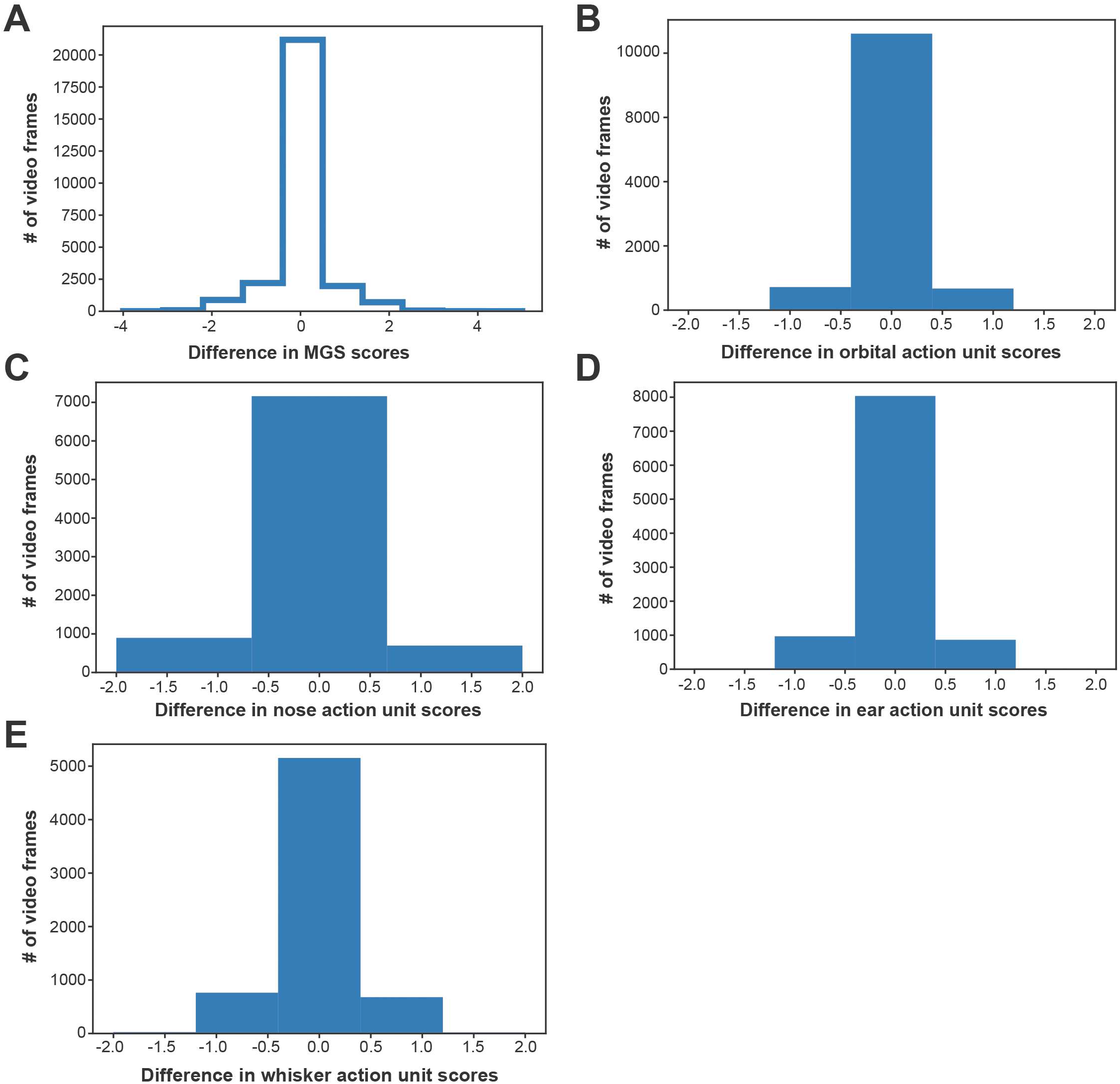
Human and MLA score differences. (A) MGS score, (B) orbital, (C) nose, (D) ear, and (E) whisker action unit scores.

**Supplementary Figure 3.**
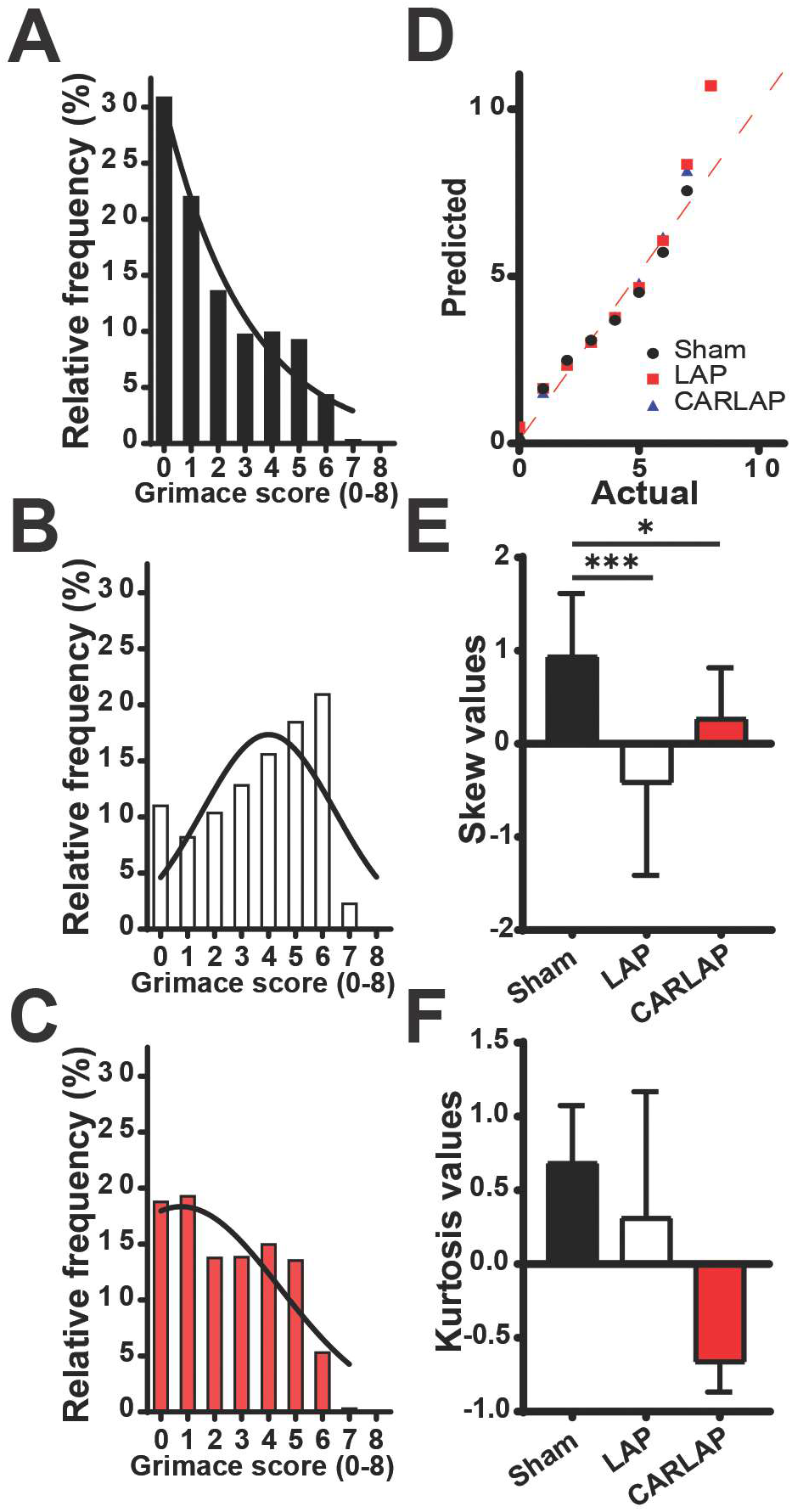
Distribution analysis of validation experiment. Frequency of each pain score for (A) sham, (B) LAP, and (C) CARLAP groups. (D) QQ plot for each experimental group. (E) Skew values for each group. (F) Kurtosis values for each group. n = 14-17 male mice/group. *p < 0.05, ***p < 0.001. Statistics: Brown-Forsythe ANOVA test.

